# Multiplex genome engineering in yeast using the TIGR-Tas system

**DOI:** 10.64898/2026.07.30.739390

**Authors:** Zhenkun Cai, Yetong Sang, Lingjie Xu, Yongfei Chang, Nga Man Wong, Jie Zhu, Chang-Ting Chen, Zehua Bao

## Abstract

Tandem Interspaced Guide RNA (TIGR)–TIGR-associated (Tas) systems are a newly discovered family of ultracompact, modular RNA-guided DNA-targeting proteins that function without a protospacer adjacent motif (PAM) requirement. Their utility as genome engineering tools in microbes remains unexplored. Here, we report the first functional implementation of TIGR-Tas in *Saccharomyces cerevisiae* for genome engineering. We show that TasR from *Parcubacteria* (ParTasR) can be programmed by user-defined tigRNAs to generate targeted DNA double-strand breaks at yeast endogenous loci. By co-delivering ParTasR with customized tigRNAs and donor templates, we achieved precise gene fragment deletion and targeted codon substitutions at multiple genomic loci. The multiplex genome engineering capability of this TIGR-Tas system was demonstrated through high-efficiency multiplex gene disruption and chromosomal assembly of a lycopene biosynthesis pathway while inactivating an endogenous gene. This work establishes TIGR-Tas as a valuable addition to the yeast genome engineering toolbox, particularly for applications requiring PAM-independent targeting or compact delivery.

## INTRODUCTION

The study of DNA double-strand break (DSB) repair in the model organism *Saccharomyces cerevisiae* laid the mechanistic foundation for precision genome engineering techniques^1–3^. Traditional genome engineering methods in yeast, relying on rare spontaneous DSB occurrence and homologous recombination (HR), have revealed the functions of genes on a genome-scale, yet remaining resource-demanding and laborious to extend across different genetic backgrounds and species^4,5^. The advent of CRISPR-Cas9 technology revolutionized the field by enabling programmable RNA-guided, site-specific DSB formation^6^. Combined with multiplex guide RNA (gRNA) expression, high-efficiency multiplex genome engineering was realized and facilitated both basic and applied research^7^. In addition, genetic screens were democratized by designing gRNA libraries targeting genome-wide^8–12^, or diversifying user-defined genomic regions^13–17^.

Despite these advances, CRISPR systems face fundamental constraints. The protospacer adjacent motif (PAM) sequence requirement restricts targetable sites: for example, SpCas9 requires a 5’-NGG PAM sequences^6,18^, limiting gRNA design flexibility particularly in AT-rich regions and constraining precise positioning of nucleotide modifications^19^. Even PAM-relaxed Cas9 variants^20,21^ cannot eliminate this requirement entirely, and comes with the compromise of reduced activity^22^. Additionally, large Cas9 proteins (∼1,368 amino acids for SpCas9) impose metabolic burden to host cells during episomal expression and limit plasmid construction and delivery, especially for multiplex genome engineering where multiple gRNAs must be co-expressed^23^. This challenge is more pronounced in non-conventional yeasts, where transformation efficiencies can be low as compared with *S. cerevisiae*^24^. Although compact Cas proteins and Cas ancestors have been described recently, their editing efficiencies are typically much lower than large Cas proteins and often require long PAM sequences, which can limit their targeting range^25^.

Recently, a mechanistically different class of RNA-guided DNA-targeting systems was discovered through computational mining of prokaryotic and viral genomes^26^. These so-called TIGR-Tas (Tandem Interspaced Guide RNA–TIGR-Associated) systems employ dual-spacer guide RNAs (tigRNAs) containing two tandem spacers that simultaneously engage both strands of the target DNA, enabling programmable DNA recognition without PAM sequence requirements. Tas proteins are remarkably compact (∼300-400 amino acids), approximately one-quarter the size of SpCas9. Two Tas monomers dimerize and form a complex with one tigRNA for DNA recognition. Among the Tas variants, TasR nucleases contain a RuvC nuclease domain at the N-terminus and generate DSBs with 8-nucleotide 3’ overhangs, with each monomer cleaving one strand of the target DNA.

Initial demonstration of TasR-mediated genome editing in human cells achieved 0.8–3.6% editing efficiency without optimization using the TasR protein from *Parcubacteria* (ParTasR)^26^, suggesting functional potential in eukaryotic systems. Given the successful adoption and improvement of CRISPR-based genome engineering in yeast^7,8,13,27^, we hypothesized that TIGR-Tas systems could be adapted for yeast genome engineering. Here, we report the development and validation of TIGR-Tas in *S. cerevisiae*. We show that TIGR-Tas is highly functional in introducing various types of genomic edits, including deletion, point mutation, and integration. TIGR-Tas also enables facile multiplex genome engineering and is highly specific. These results establish TIGR-Tas as a valuable addition to the yeast genome engineering toolbox, particularly for applications requiring PAM-independent targeting or compact delivery.

## RESULTS

### Validation of in vivo DNA cleavage by TIGR-Tas

We chose ParTasR for testing in *S. cerevisiae* considering its higher editing efficiency in human cells as compared to other Tas variants^26^. Two components, the ParTasR protein and the tigRNA are required for DNA targeting (**Fig. 1A**). The tigRNA consists of 36 nucleotides containing two 9-nucleotide spacers (spacer A and spacer B) flanked by the conserved box C (CCA) and box D (UG) motifs. These spacers jointly specify the target for the TasR protein, with spacer A pairing to one strand of the target DNA and spacer B pairing to the other strand. To express both components in yeast cells, we constructed a single plasmid expressing a human codon-optimized ParTasR gene driven by the constitutive TEF1 promoter and a tigRNA driven by the type Ⅲ SNR52 promoter, which is commonly used for expressing CRISPR RNAs (**Fig. 1B** and **Supplementary Fig. 1**). ParTasR was fused with nuclear localization signals to facilitate nuclear entry after expression. To assess whether ParTasR exhibits programmable nuclease activity in *S. cerevisiae*, we adapted a previously established yeast-based cleavage assay used to evolve CjCas9 and TnpB (**Fig. 1C**)^28,29^. A target sequence harboring a premature stop codon was inserted between two 100 bp repeats within the genomic *ADE2* locus, disrupting the gene and making the cell auxotrophic for adenine (**Fig. 1D**). A functional *ADE2* gene can be restored through single-strand annealing (SSA)-mediated repair following a targeted double-strand break. We designed four tigRNAs targeting distinct sites between the *ADE2* duplication regions (**Fig. 1E**). The only design criteria we adopted were the exclusion of more than 4 consecutive T’s in the tigRNA sequence to avoid early transcription termination and the uniqueness of the 18 bp target site within the yeast genome to minimize potential off-target effects. Among the four tigRNAs, three conferred significant survival rates of the transformed yeast cells on adenine-deficient medium, while one showed no detectable activity (**Fig. 1F** and **Supplementary Fig. 2**). These results indicated that TIGR-Tas was functionally expressed in our system and mediated efficient RNA-guided DNA cleavage in *S. cerevisiae*.

**Figure 1.**
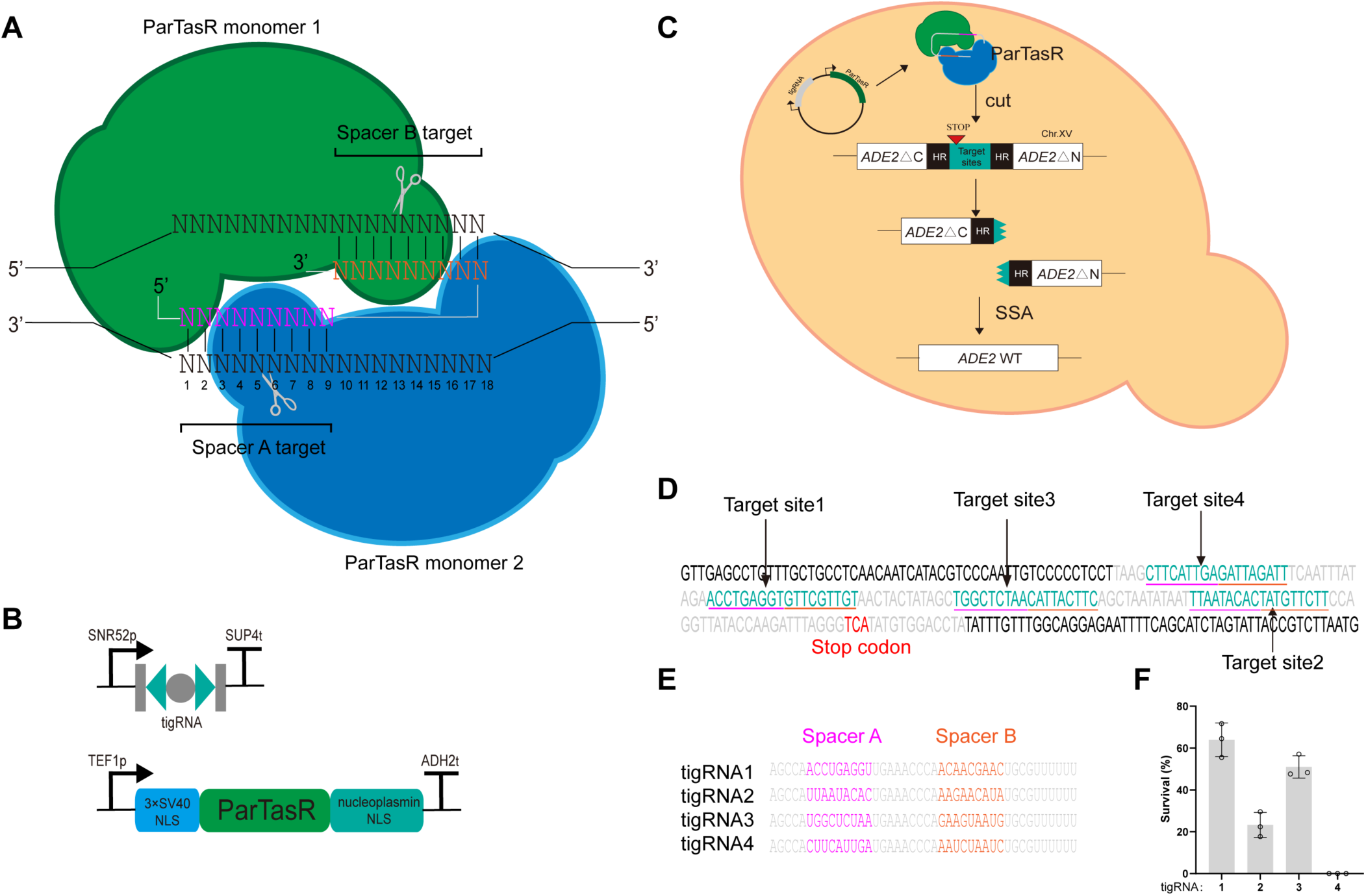
TIGR-Tas system design and validation in *S. cerevisiae*. (**A**) Illustration of ParTasR dimer complexed with a tigRNA to direct endonuclease activity toward the genomic DNA target. The tigRNA contains two spacer sequences separated by conserved repeat elements, with one spacer pairing to one strand of the target DNA and the other spacer pairing to the opposite strand. (**B**) Design of the tigRNA and ParTasR expression constructs. tigRNA expression was driven by the type Ⅲ SNR52 promoter and terminated by the SUP4 3’ flanking sequence. ParTasR was fused with three tandem SV40 nuclear localization signals (NLS) at the N-terminus and an additional NLS at the C-terminus and was expressed under the constitutive TEF1 promoter. (**C**) Schematic of the single-strand annealing (SSA) repair assay. Target sites to be tested were inserted between two repeat regions of the engineered genomic *ADE2* locus. Upon TasR-mediated cleavage at the target site, SSA repair restores *ADE2* expression, enabling cell survival on adenine-deficient medium. (**D**) Schematic of the *ADE2* reporter locus showing the detailed sequences of the repeats (black) and the target sites (cyan). Spacer A target and spacer B target were annotated with magenta and orange underlines. The reverse-complemented premature stop codon was annotated in red. (**E**) Full sequences of the four tigRNAs. Each tigRNA is 36-nucleotide long containing two 9-nucleotide spacers (spacer A and spacer B) that together recognize the inserted target site. Conserved sequences are denoted in grey. (**F**) Survival rates of yeast cells transformed with plasmids expressing ParTasR and each of the four tigRNAs on adenine-deficient medium. Data represents mean ± SD from three independent experiments.

### Precise gene fragment deletion mediated by TIGR-Tas

To further evaluate whether ParTasR could mediate precise chromosomal edits given its high DNA cleavage activity as reported via the SSA assay, we designed another four tigRNAs targeting the endogenous undisrupted *ADE2* sequence and co-delivered a repair template with 100 bp homology arms to enable precise fragment deletion defined by the arms (**Fig. 2A**). The four tigRNA targets, designated as ADE2.a-d, all reside in between the two arms. All components—including ParTasR driven by TEF1p, tigRNA expressed by SNR52p, and the donor repair template—were assembled into a single plasmid. This all-in-one design allows for the sustained presence of the plasmid-harbored donor and potentially increases editing efficiency, as compared with linear repair donors^14^. Among the four tigRNAs tested, three showed no detectable activity, while one exhibited robust editing activity with over 60% editing efficiencies as measured by the percentage of yeast colonies turning pink (**Fig. 2B**). Sequencing of the pink colonies further confirmed the precise fragment deletion as programmed by the donor at the target site (**Supplementary Fig. 3**). This result further indicated that, with an efficient tigRNA, the TIGR-Tas system was able to target endogenous sequences and perform genome engineering through homologous recombination.

**Figure 2.**
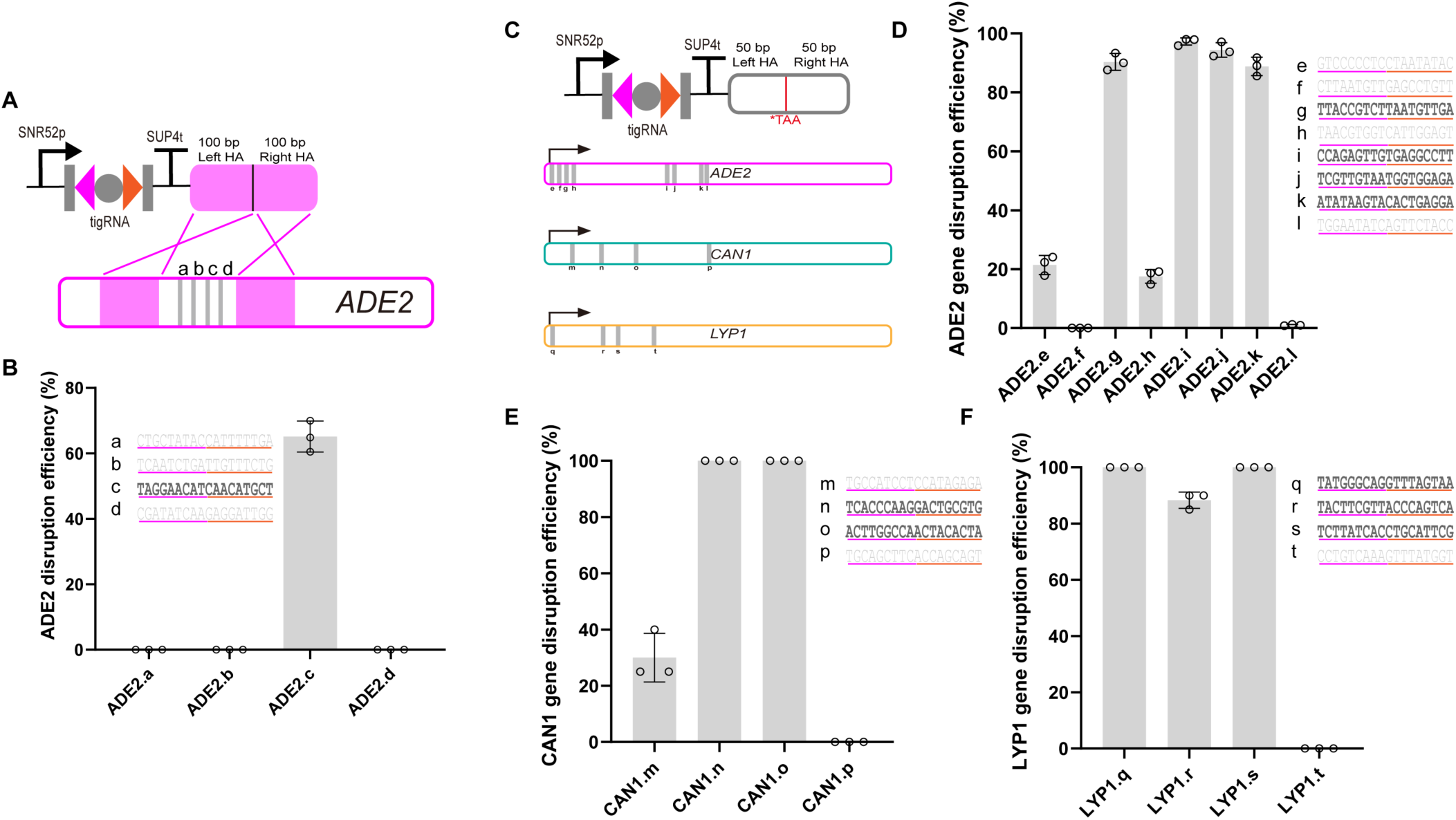
Precise genome editing at endogenous loci. **(A)** Schematic of the experimental design for targeted 200 bp deletion at the *ADE2* locus. The repair donor template, containing two 100 bp homology arms flanking the deletion region, was placed immediately downstream of the tigRNA expression cassette on the same plasmid. Upon TasR-mediated cleavage, homology-directed repair (HDR) using the donor template results in the precise deletion of the 200 bp fragment. **(B)** Deletion efficiencies of four tigRNAs targeting different sites within the wild-type *ADE2* locus. Editing efficiencies were determined by calculating the proportion of pink colonies among total colonies and validated by Sanger sequencing of selected colonies. Efficient target sites are highlighted in bold. Data represents mean ± SD from three independent experiments. **(C)** Schematic of the experimental design for precise codon substitution at each target locus. The repair template, containing two 50-bp homology arms flanking the target site, was placed immediately downstream of the tigRNA expression cassette on the same plasmid. A TAA stop codon along with synonymous mutations was introduced within the spacer-targeting region, which prevents re-cleavage after editing. Eight tigRNAs were selected for *ADE2*, and four tigRNAs each for *CAN1* and *LYP1*. All tigRNAs target the first half of the target gene so that the protein product will be inactivated. **(D)** Editing efficiencies of eight tigRNAs targeting *ADE2*. **(E)** Editing efficiencies of four tigRNAs targeting *CAN1*. **(F)** Editing efficiencies of four tigRNAs targeting *LYP1*. All positive clones were further validated by Sanger sequencing. Efficient target sites are highlighted in bold. Data represents mean ± SD from three independent experiments.

### Precise codon substitutions mediated by TIGR-Tas

We next tested the ability of ParTasR to mediate precise codon substitutions in the yeast genome by designing repair templates containing target codon substitution to a stop codon. Within the spacer-targeting region of the template, we introduced a TAA stop codon mutation along with synonymous mutations to disrupt the seed region critical for tigRNA recognition, as reported previously^26^, thereby preventing cleavage of the template as well as the edited sequence after homologous recombination. The repair template consists of approximately 50 bp homology arms (**Fig. 2C**). To assess the overall likelihood of finding efficient tigRNAs, eight additional tigRNAs were designed to target randomly selected positions within the first half of the *ADE2* gene (designated as ADE2.e-l, **Fig. 2C**), with donor templates incorporated into the same all-in-one plasmid. The positioning of the target sites within the first half of the gene ensures that once edited, the mutated gene sequence will result in a phenotypic readout due to early truncation of the translation product. Phenotypic analysis revealed that four of the eight tigRNAs supported highly efficient editing (all over 80%, **Fig. 2D**), while the other four tigRNAs showed less than 20% editing efficiencies or no editing at all. Sequencing confirmed precise codon substitutions as programmed by corresponding donors at all active target sites (**Supplementary Fig. 4**). To assess the generalization of TIGR-Tas editing across different genomic loci, we similarly tested four tigRNAs each at the *CAN1* and *LYP1* loci and, in each case, identified half or more highly active tigRNAs with near 100% editing efficiencies (**Fig. 2E,F**).

### Dual-locus editing mediated by TIGR-Tas

Having demonstrated efficient single locus editing, we set out to evaluate the multiplex genome editing capability of the TIGR-Tas systems, as multiplex genome editing is a critical advanced feature of genome engineering technologies to facilitate yeast strain engineering for synthetic biology. We first performed dual-locus editing within a single gene (**Fig. 3A**). To do this, we selected two highly active tigRNAs targeting two distinct sites (ADE2.g and ADE2.i, **Fig. 2D**) in the *ADE2* gene, along with their corresponding repair templates, and implemented three different multiplexing strategies to express the two tigRNAs from the same plasmid. In the first strategy (dual promoter), the two tigRNAs were expressed independently, each driven by its own promoter: one by the SNR52 promoter and the other by the tRNA^Tyr^ promoter^30,31^. In the second strategy (tigRNA-tRNA array), the two tigRNAs were assembled into a tandem array separated by a tRNA^Gly^ sequence, which will be post-transcriptionally cleaved by endogenous enzymes to release the two tigRNAs^32^. In the third strategy (TIGR array), the two tigRNAs were placed in tandem without any intervening sequences (**Fig. 3A**). For all strategies, the repair templates were placed in tandem downstream of the tigRNA expression cassettes. Upon plating right after transformation, all three strategies yielded over 90% pink colonies, indicating efficient *ADE2* disruption. However, Sanger sequencing of pink colonies revealed a key difference: the dual-promoter and TIGR array strategies achieved successful dual-locus editing at both ADE2.g and ADE2.i loci in over 90% of the colonies tested (representative sequencing results for the TIGR array strategy was shown in **Fig. 3B**). In contrast, the tigRNA-tRNA array strategy, despite showing a high proportion of pink colonies, only yielded editing at the ADE2.g locus, with no detectable editing at the ADE2.i locus. We observed that when transformants from the tigRNA-tRNA array strategy were first cultivated in liquid SC selective medium for 3 days before plating, the dual-locus editing efficiency also reached over 90% (**Fig. 3C-E**). These results demonstrate that all three multiplexing strategies are effective for dual-locus editing, with the dual-promoter and TIGR array strategies enabling faster dual editing, while the tigRNA-tRNA array strategy requires an additional liquid cultivation step to achieve comparable performance.

**Figure 3.**
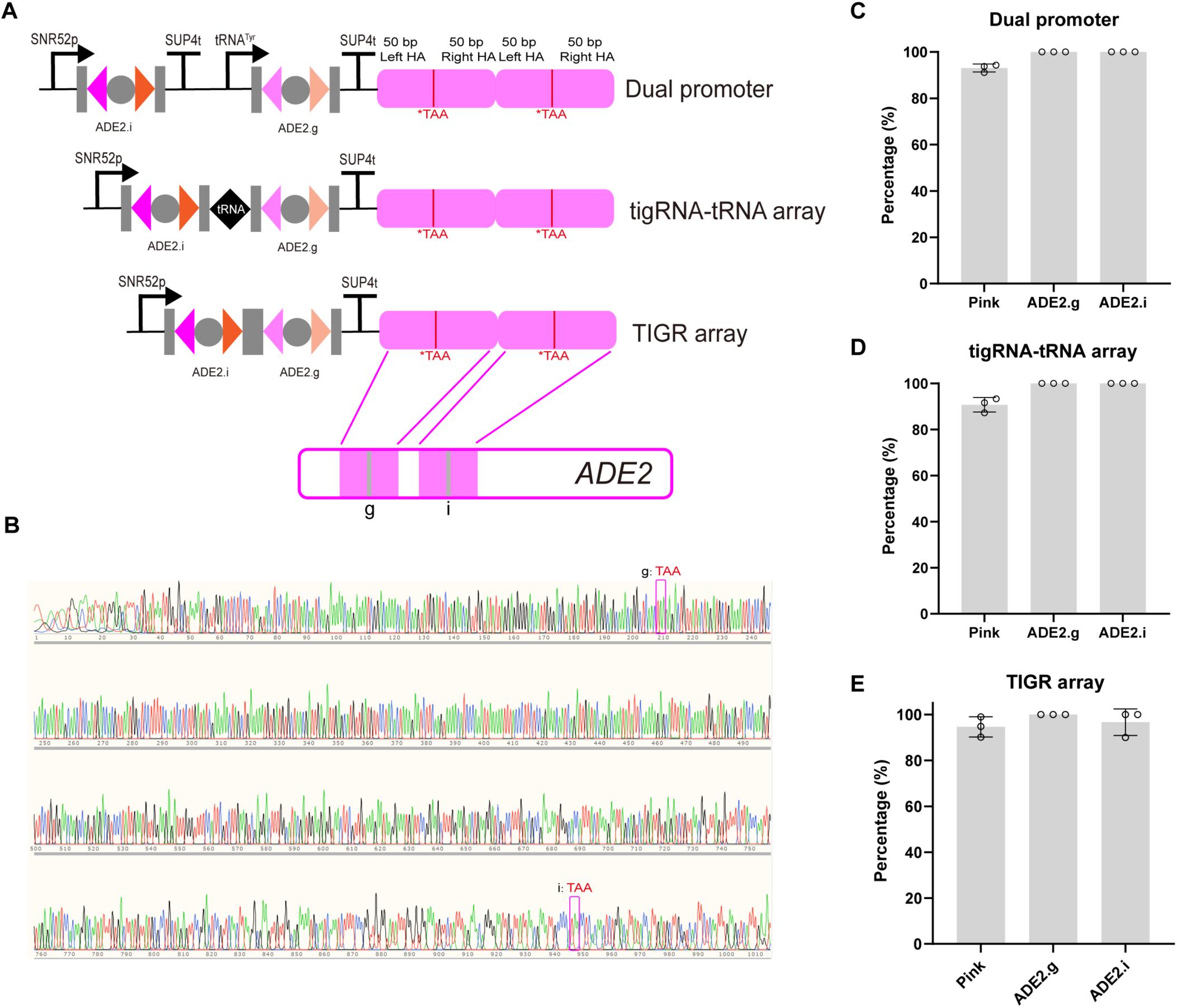
Dual-locus editing of *ADE2* using three multiplexing strategies. **(A)** Schematic of the three strategies for dual-locus editing of *ADE2*. In the dual-promoter strategy, two tigRNAs are expressed from separate expression cassettes, each driven by its own promoter (SNR52 promoter and tRNA^Tyr^ promoter). In the tigRNA-tRNA array strategy, two tigRNAs are assembled into a tandem array separated by a tRNA (tRNA^Gly^), allowing processing by endogenous tRNA-processing enzymes to release individual tigRNAs from a single transcript. In the TIGR array strategy, two tigRNAs are placed in tandem without any intervening sequences. **(B)** A representative Sanger sequencing chromatogram showing the introduction of TAA stop codons at both target sites within the *ADE2* gene (shown for the TIGR array strategy). **(C)** Dual-locus editing efficiencies using the dual-promoter strategy. The proportion of pink colonies indicative of *ADE2* disruption is shown. From the pink colonies, 20 were randomly selected and subjected to Sanger sequencing to determine the introduction efficiency of the TAA stop codon at each of the two target sites. **(D)** Dual-locus editing efficiencies using the tigRNA-tRNA array strategy. The proportion of pink colonies and the TAA introduction efficiency at both target sites are shown. (**E)** Dual-locus editing efficiencies using the TIGR array strategy. The proportion of pink colonies and the TAA introduction efficiency at both target sites are shown. Data represents mean ± SD from three independent experiments.

### Triple-gene editing mediated by TIGR-Tas

Next, we evaluated multiplex editing of distant genes across the yeast genome, which is required for various strain engineering tasks. We selected highly active tigRNAs targeting the ADE2.i, CAN1.q, and LYP1.n sites (**Fig. 2**), each paired with its corresponding repair template. Akin to dual-locus editing, three strategies were implemented. In the first strategy (triple promoter), the three tigRNAs were expressed independently, each driven by its own promoter: the SNR52 promoter, the tRNA^Tyr^ promoter, and the tRNA^Pro^ promoter^30,31^. In the second strategy (tigRNA-tRNA array), the three tigRNAs were assembled into a tandem tigRNA-tRNA array, separated by different tRNA sequences^33^. The use of distinct tRNA sequences between tigRNAs was intended to minimize sequence homology, thereby facilitating plasmid construction and reducing the risk of homologous recombination that could lead to plasmid rearrangement and loss of tigRNAs. In the third strategy (TIGR array), the three tigRNAs were placed in tandem without any intervening sequences (**Fig. 4A**). Upon plating right after transformation, the three strategies showed markedly different outcomes. The triple-promoter strategy yielded almost entirely pink colonies and the TIGR array yielded mostly pink colonies, while the tigRNA-tRNA array (with tRNA^Gly^-tRNA^Thr^) produced the least frequent pink colonies and sectored pink-white chimeric colonies (**Supplementary Fig. 5A**), indicating progressively slower editing kinetics. For accurate efficiency measurement, transformants from all three strategies were first plated, then scraped, cultivated in liquid SC medium for 3 days, and re-plated. After liquid cultivation, the triple-promoter strategy achieved the highest triple-editing efficiency with over 80% (**Fig. 4B**). The tigRNA-tRNA array showed lower but detectable triple-editing efficiency (**Fig. 4C**). The TIGR array also achieved substantial triple-editing efficiency, with the advantage of a compact array structure where tigRNAs are placed in tandem without any intervening sequences (**Fig. 4D**). From the dual-locus and triple-gene editing results of the tigRNA-tRNA array strategy, we suspect that the lower efficiencies at the *ADE2* locus may be attributable to the specific tRNA used. To improve *ADE2* editing efficiency in the tigRNA-tRNA array strategy, we replaced tRNA^Gly^ with tRNA^Ser^ or tRNA^Leu^. However, neither tRNA^Ser-^tRNA^Thr^ nor tRNA^Leu^-tRNA^Thr^ significantly increased the proportion of pink colonies (**Supplementary Fig. 5B**). Nonetheless, these results establish that all three multiplexing strategies are compatible with TIGR-Tas for multiplex genome engineering in yeast, with the TIGR array strategy offering a favorable balance between high efficiency and compact design.

**Figure 4.**
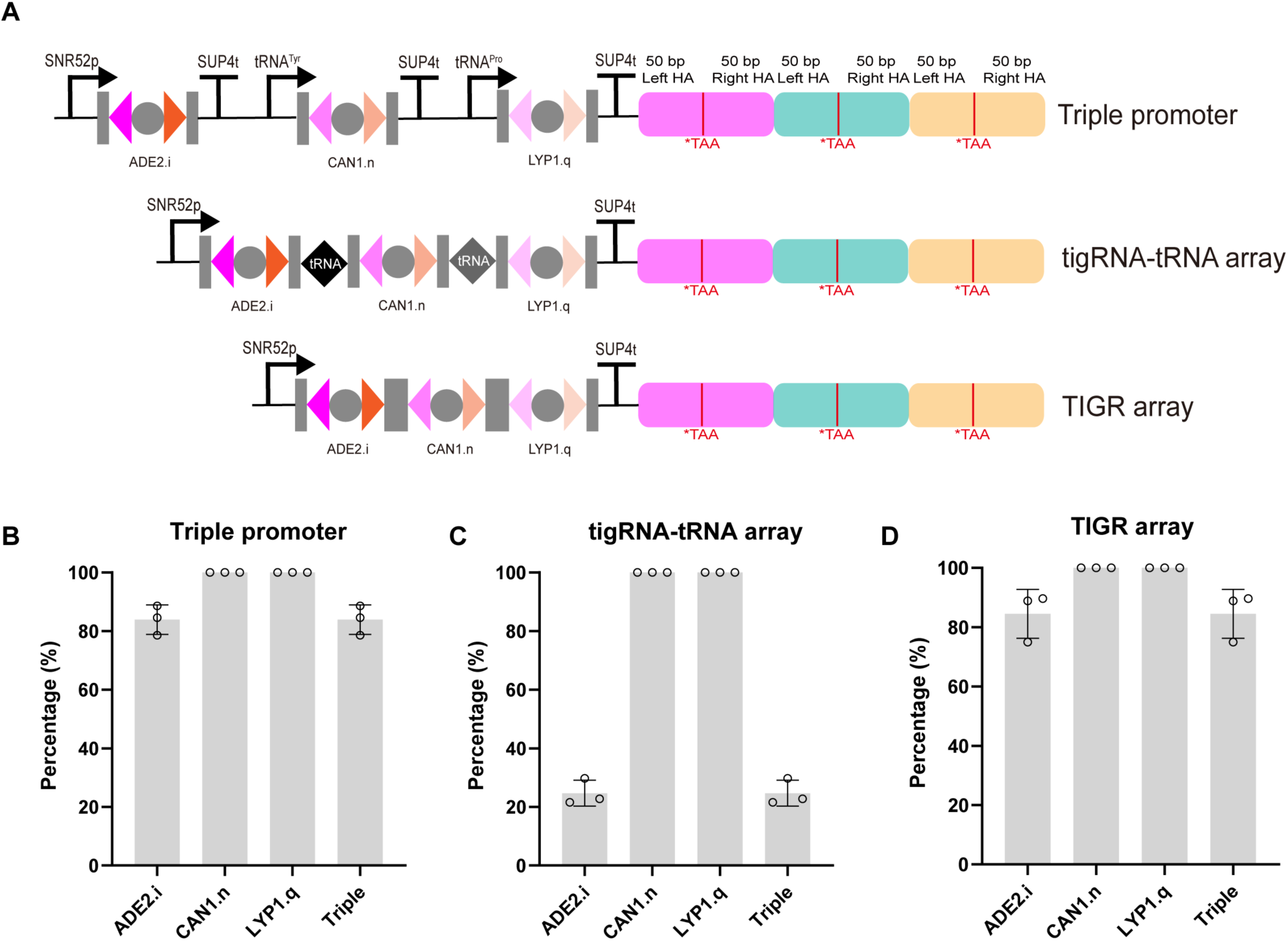
Triple-gene editing using three multiplexing strategies. **(A)** Schematic of the three strategies for triple-gene editing targeting *ADE2*, *CAN1*, and *LYP1*. In the triple-promoter strategy, three tigRNAs are expressed from separate expression cassettes, each driven by an independent promoter (SNR52 promoter, tRNA^Tyr^ promoter, and tRNA^Pro^ promoter). In the tigRNA-tRNA array strategy, three tigRNAs are assembled into a tandem array separated by tRNAs (tRNA^Gly^ and tRNA^Thr^), allowing processing by endogenous tRNA-processing enzymes to release individual tigRNAs from a single transcript. In the TIGR array strategy, three tigRNAs are placed in tandem without any intervening sequences. **(B)** Triple-gene editing efficiencies using the triple-promoter strategy. From left to right, the four bars represent: (i) the proportion of pink colonies indicative of *ADE2* disruption; (ii) the proportion of pink colonies that survive L-canavanine selection (indicative of *CAN1* disruption); (iii) the proportion of pink colonies that survive thialysine selection (indicative of *LYP1* disruption); and (iv) the triple-gene editing efficiency, defined as the proportion of colonies that are pink and both canavanine and thialysine resistant. **(C)**, Triple-gene editing efficiencies using the tigRNA-tRNA array strategy. The four bars represent the same metrics as in B. **(D)** Triple-gene editing efficiencies using the TIGR array strategy. The bars represent the same metrics as in B. All positive clones were further validated by Sanger sequencing. Data represents mean ± SD from three independent experiments.

### Metabolic pathway engineering using TIGR-Tas

To evaluate the potential of TIGR-Tas for metabolic engineering applications, we performed targeted gene integration using the TIGR-Tas system. First, we used four ADE2-targeting tigRNAs (ADE2.g, ADE2.i, ADE2.j, ADE2.k) to integrate a linear donor fragment containing a TDH3p-*EGFP*-ADH1t expression cassette into the yeast genome. As a control, we also performed transformations with TIGR-Tas alone (without the linear donor fragment) to assess indel efficiency induced by ParTasR cleavage (**Fig. 5A**). Notably, pink colonies were observed without a repair template, with the proportion of pink colonies (reflecting indel efficiency) ranging from 2.83% to 33.33% depending on the tigRNA (**Fig. 5B**). Sanger sequencing of the four endogenous loci confirmed random indels at the target sites (**Supplementary Fig. 6**), demonstrating that ParTasR can achieve gene disruption through the non-homologous end joining (NHEJ) DNA repair pathway. This suggests that the TIGR-Tas system may be extended to other fungi hosts with high NHEJ efficiency, such as *Yarrowia lipolytica*^34^ and *Komagataella phaffii*^35^. In the presence of the EGFP donor, three tigRNAs (ADE2.g, ADE2.i, ADE2.k) successfully mediated integration of the EGFP expression cassette, achieving EGFP-positive rates ranging from 31.4% to 81.1%. In contrast, ADE2.j showed low indel efficiency and low EGFP-positive rate, indicating weaker cleavage activity (**Fig. 5B**). Next, we tested the ability of TIGR-Tas to mediate genomic integration of a more complex metabolic pathway. We chose the three-gene lycopene biosynthesis pathway (*crtE*, *crtYB*, *crtI*) as a test case, using a highly active *CAN1*-targeting tigRNA for integration (**Fig. 5C**). Successful integration resulted in red colonies due to lycopene accumulation (**Fig. 5D**). The integration efficiency for the three-gene pathway reached 11.4-35.3%, which is lower compared to single-gene integration or single codon substitutions, likely due to the longer pathway length and the increased recombination events required for in vivo pathway assembly^36^. The above results demonstrate that the TIGR-Tas system can efficiently mediate both single-gene expression cassette integration and multi-gene metabolic pathway assembly, showing potential for metabolic engineering and synthetic biology applications.

**Figure 5.**
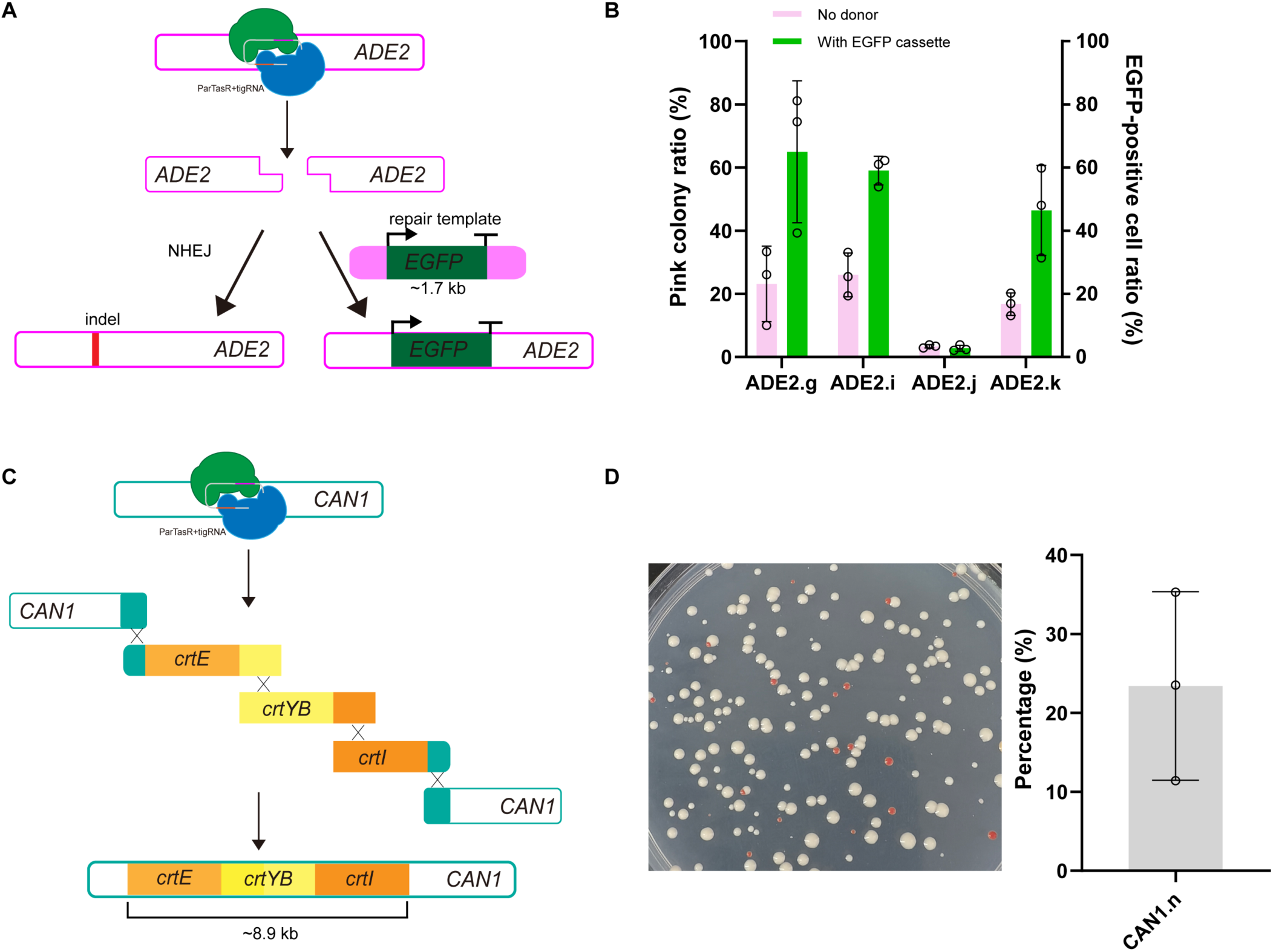
Metabolic pathway engineering using TIGR-Tas in *S. cerevisiae*. **(A)** Schematic of the experimental design for *EGFP* integration. A linear donor fragment containing the TDH3p-*EGFP*-ADH1t expression cassette flanked by 50 bp homology arms was co-delivered with ParTasR and an *ADE2*-targeting tigRNA. The control (without donor) was used to assess the indel efficiency induced by ParTasR cleavage. **(B)** EGFP-positive rates mediated by four *ADE2*-targeting tigRNAs (ADE2.g, ADE2.i, ADE2.j, ADE2.k) with or without the EGFP donor. Data represent mean ± SD from three independent experiments. **(C)** Schematic of the three-gene lycopene biosynthesis pathway integration. Three linear donor fragments containing ENO2p-crtE-ENO2t, TEF2p-crtYB-TEF2t, and CDC19p-crtI-CDC19t were assembled in vivo with 150 bp overlapping regions and 50 bp flanking homology arms targeting the *CAN1* locus. **(D)** Representative plate showing red colonies after lycopene pathway integration and integration efficiencies of the three-gene lycopene pathway. Red color indicates successful integration and lycopene production. Data represent mean ± SD from three independent experiments.

### Specificity of TIGR-Tas targeting

DNA targeting specificity is a critical feature of all genome engineering tools to avoid off-target effects. To assess the specificity of TIGR-Tas editing, we designed a mismatch tolerance assay using two highly active tigRNAs targeting *ADE2* (ADE2.i and ADE2.k). For each tigRNA, we introduced single-nucleotide mismatches at each of the 18 positions within the combined spacer A and spacer B sequences (9 nt each), substituting the original base with its complementary transversion (A↔T, C↔G). A total of 36 mutant tigRNAs (18 for each tigRNA) were generated, and the original repair templates introducing the TAA stop codon were retained. Each mutant tigRNA was co-expressed with ParTasR and the repair donor from the same all-in-one plasmid. After transformation, editing efficiency was determined by calculating the proportion of pink colonies, as described above. If the mismatch did not affect TIGR-Tas cleavage activity, the repair template would introduce the TAA stop codon via homologous recombination, resulting in pink colonies. Conversely, if the mismatch abolished cleavage activity, the stop codon could not be introduced, and colonies remained white (**Fig. 6A**). Remarkably, the vast majority of single mismatches almost completely abolished editing activity (**Fig. 6B, C**). For both ADE2.i and ADE2.k, only mismatches at positions near the 5’ or 3’ edges of the spacer regions retained detectable activity, while mismatches at internal positions eliminated editing entirely. We further tested selected double mismatches combining two tolerant edge mismatches. Such double mismatches completely abolished editing activity (**Fig. 6B, C**). The high mismatch sensitivity observed is likely attributed to the dual-spacer targeting mechanism of the TIGR-Tas system, where each spacer contains a seed region that is highly sensitive to mismatches. Based on these results, we propose that tigRNAs should be designed to ensure that no perfectly matching site exists in the yeast genome for the 18 bp targeting sequence, and that any potential off-target site differs from the on-target sequence by at least 2 mismatches to minimize the risk of off-target editing.

**Figure 6.**
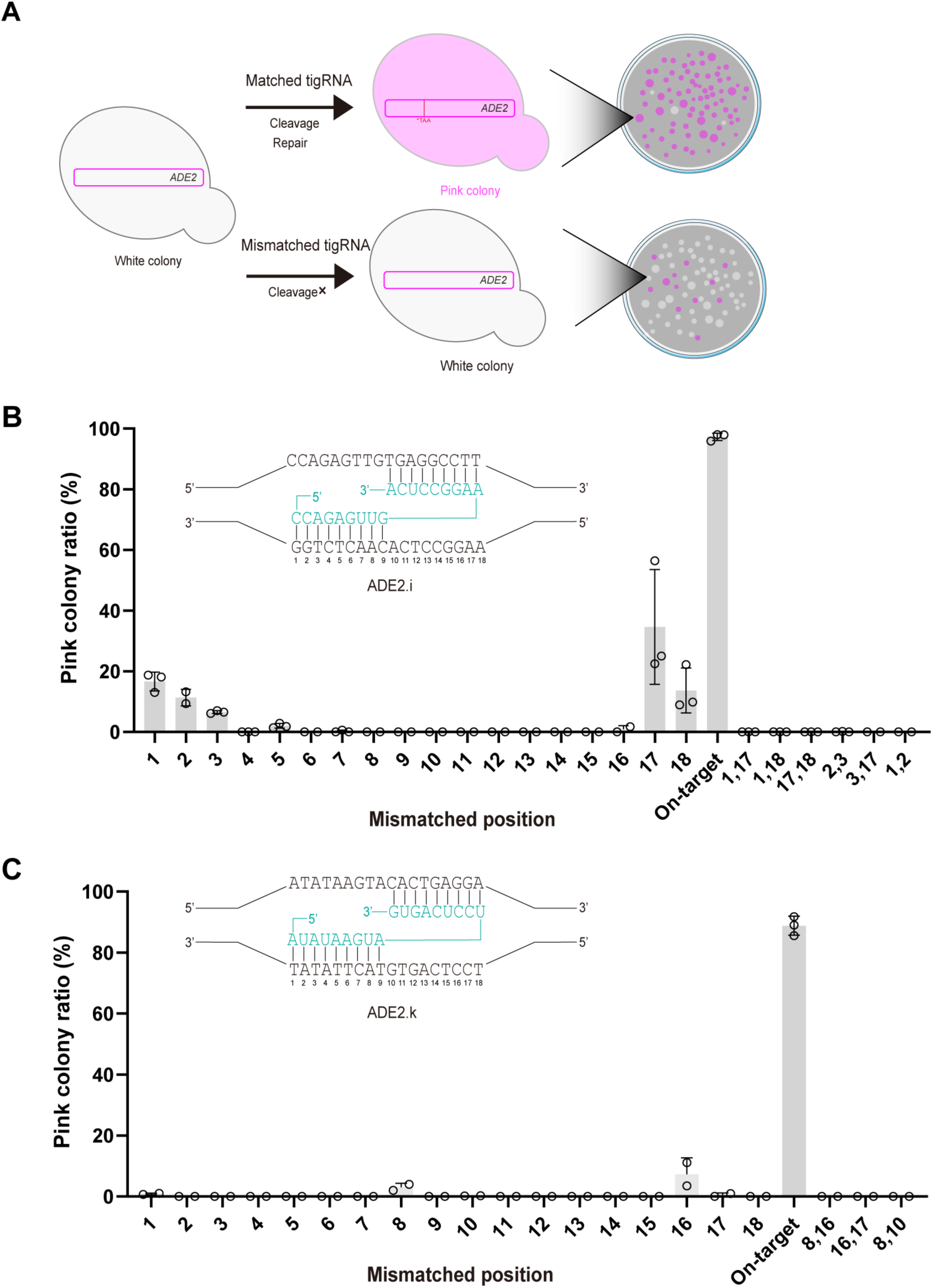
Specificity of TIGR-Tas targeting. **(A)** Schematic of the mismatch tolerance assay. When the tigRNA is perfectly matched to the target, ParTasR-mediated cleavage stimulates homologous recombination with the donor template containing the TAA stop codon, resulting in pink colonies. When the tigRNA contains mismatches that impair cleavage, no TAA stop codon is introduced, and colonies remain white. Perfectly matched tigRNA yields pink colonies. Mismatched tigRNA yields white colonies or only a few pink colonies depending on the mismatch position. **(B)** Mismatch tolerance profiles for ADE2.i tigRNA. Editing efficiencies for each of the 18 single-nucleotide mismatch positions are shown. The positions of mismatches are numbered from 1 to 18 along the target sequence. Selected double mismatches combining two tolerant edge mismatches are also indicated. **(C)** Mismatch tolerance profiles for ADE2.k tigRNA. Selected double mismatches combining two tolerant edge mismatches are also indicated. Data represent mean ± SD from at least two independent experiments.

## DISCUSSION

In this study, we demonstrated that the TIGR-Tas system—a recently discovered family of RNA-guided DNA-targeting enzymes distinct from CRISPR-Cas—is functional in *S. cerevisiae* for genome engineering. We showed that ParTasR, when expressed in yeast, exhibited programmable nuclease activity at multiple endogenous loci, enabling both precise gene fragment deletions and codon substitutions. Furthermore, we demonstrated that TIGR-Tas can efficiently mediate both single-gene expression cassette integration and multi-gene metabolic pathway assembly, highlighting its potential for genomics and metabolic engineering applications.

Currently, tigRNA design lacks well-defined sequence-activity rules, in contrast to CRISPR systems where extensive empirical data have enabled the development of reliable gRNA design tools. In this study, our design strategy was limited to avoiding poly-T tracts (≥4 thymidines) and selecting sequences with high predicted genomic specificity. Despite this minimal guidance, we successfully identified highly active tigRNAs targeting *ADE2*, *CAN1*, and *LYP1* through random selection, with approximately half of the tested tigRNAs exhibiting high activity—a success rate comparable to that of CRISPR-Cas9. Using a mismatch tolerance assay, we further found that the vast majority of single mismatches within the 18 bp spacer sequences almost completely abolished editing activity, and double mismatches combining two tolerant edge mismatches eliminated the residual activity associated with the tolerant single mismatches. These results suggest that the specificity of TIGR-Tas editing can be ensured by carefully selecting the tigRNA targeting sequence. However, the molecular determinants of tigRNA efficiency remain unclear. A systematic profiling of tigRNA sequence-activity relationships using high throughput libraries in future studies could provide the data needed to develop predictive models and rational design rules, analogous to the gRNA activity prediction approaches established for CRISPR systems in yeasts^37^.

One of the most distinctive features of TIGR-Tas systems is their lack of a PAM requirement, which expands the accessible genome space, particularly in AT-rich regions such as promoter elements. Furthermore, whereas CRISPR-Cas9 gRNAs require a conserved scaffold sequence for Cas protein binding, tigRNAs functioned without such a bulky scaffold. This scaffold-free architecture enables tigRNAs to be processed from a single transcript, similar to the crRNA processing mechanism of Cas12a^38^. We developed and compared three multiplex editing strategies: a multiple-promoter approach, a tRNA-interspaced array, and a compact TIGR-array where tigRNAs are placed in tandem without intervening sequences. Remarkably, the TIGR-array alone was sufficient to process multiple tigRNAs from a single transcript and achieved multiplex editing efficiencies comparable to the multiple-promoter approach, while substantially outperforming the tRNA-interspaced array. For triple-gene editing, the triple-promoter strategy achieved the highest efficiency, followed by the TIGR-array, while the tRNA-interspaced array showed lower efficiency. The TIGR-array, with its compact design and efficient processing, offers a streamlined platform for multiplex genome engineering. As demonstrated in this study, using the TIGR-array, we achieved simultaneous editing of three genes from a single all-in-one plasmid, suggesting that TIGR-Tas is particularly well-suited for highly multiplex applications. This capability parallels recent advances in orthogonal CAST systems that enabled parallel multi-locus editing in a single step^39^.

TasR is one member of a larger family of TIGR-associated proteins that exhibit diverse domain architectures, including TasA (no nuclease domain fusion) and TasH (HNH fusion). In this study, we focused on TasR from *Parcubacteria* (ParTasR), which contains a RuvC nuclease domain and cleaves DNA to generate 8-nucleotide 3’ overhangs. However, the suitability of other family members for yeast genome manipulation remains to be further explored. It remains to be determined whether TasA proteins can function as RNA-guided DNA-binding platforms for transcriptional regulation or epigenetic modification, and whether TasH nucleases possess comparable editing efficiencies and offer distinct features than TasR. The modular architecture of TIGR systems—where the core dimerization domain and tigRNA binding domain can be fused to different effector domains—suggests that this family may provide a rich source of programmable tools beyond nuclease activity. Exploring the functional diversity of TIGR-associated proteins could unlock new capabilities for synthetic biology, potentially enabling applications ranging from targeted transcriptional activation to epigenetic reprogramming, as has been achieved with dCas9-based platforms^40,41^. The compact nature of TIGR-Tas systems could be further leveraged by fusing them with compact TADs, such as those recently reported^42^.

The robust activity of TasR in yeast, combined with its compact size, PAM-free targeting, and scaffold-free tigRNA architecture, positions TIGR-Tas as a promising platform for metabolic engineering. Future work could couple TIGR-Tas-mediated multiplex integration with other engineering approaches to create more efficient synthetic pathways and cell factories^43^. The modular architecture of TIGR-Tas—where tigRNAs serve as programmable DNA recognition modules and Tas proteins as functional effectors—aligns with a broader paradigm in synthetic biology: the development of modular, extensible toolkits for programmable gene control. This paradigm has been successfully exemplified by systematic libraries of chromatin regulators fused to programmable DNA-binding domains for functional interrogation^44^, and by orthogonal hormone-inducible systems for multigene expression control in cellular engineering applications^45^. As more TIGR-Tas systems are characterized and their sequence-activity relationships elucidated, we anticipate that this family will provide a valuable addition to the synthetic biology toolkit for yeast, enabling applications that are challenging with current technologies.

## METHODS

### Strains and media

The *S. cerevisiae* strain BY4741 (MATa *his3Δ0 leu2Δ0 met15Δ0 ura3Δ0*) was used as the parent strain for all TIGR-Tas experiments. From this parent, we generated an *ADE2* reporter strain for DNA cleavage characterization by integrating a target sequence flanked by 100 bp repeats into the genomic *ADE2* locus, disrupting the open reading frame at the same time. Yeast cells were routinely grown in yeast extract peptone dextrose (YPD) medium at 30 ℃ with shaking at 220 rpm. Transformants were selected on synthetic complete (SC) medium lacking appropriate auxotrophic supplements. For *ADE2* disruption restoration, cells were plated on SC-adenine plates. For *ADE2* knockout, colonies were scored for pink/white phenotype on SC-U plates supplemented with limited adenine (10 μg/mL). For *CAN1* and *LYP1* editing, colonies were randomly selected from SC-U plates and replica-plated onto SC-arginine plates containing 60 μg/mL L-canavanine (Sangon Biotech, Shanghai, China) and SC-lysine plates containing 250 μg/mL thialysine (S-2-aminoethyl-L-cysteine, Sigma-Aldrich, St. Louis, MO, USA), respectively, to determine editing efficiencies.

### Plasmid construction

The ParTasR coding sequence was codon-optimized for expression in human cells and commercially synthesized (SynbioB, Tianjin, China). The amino acid sequence of ParTasR is provided in **Supplementary Table 1**. The ParTasR gene fragment was fused with the constitutive TEF1 promoter and the ADH2 terminator into a single expression cassette by overlap extension PCR, with three tandem SV40 nuclear localization signals (NLS) appended to the N-terminus and an additional nucleoplasmin NLS added to the C-terminus. The ParTasR expression cassette was assembled into the pCRCT backbone (Addgene plasmid #60621) via Gibson Assembly (Catalog # E2611, New England Biolabs, Ipswich, MA, USA), generating the pCRCT-ParTasR plasmid. For single-gene editing, tigRNA-donor fragments were commercially synthesized (SynbioB, Tianjin, China) and assembled into the pCRCT-ParTasR plasmid by Gibson Assembly. For the multiple-promoter strategy, the tRNA^Tyr^ and tRNA^Pro^ promoters were amplified from *S. cerevisiae* genomic DNA using appropriate primers (**Supplementary Table 2**) and assembled into the pCRCT-ParTasR plasmid together with corresponding repair donors via Gibson Assembly. The RNA polymerase Ⅲ promoter sequences used to express tigRNAs are listed in **Supplementary Table 3**. For the tigRNA-tRNA array strategy, the entire tigRNA-tRNA-donor array was commercially synthesized (SynbioB, Tianjin, China) and assembled into the pCRCT-ParTasR plasmid by Gibson Assembly. For the TIGR-array strategy, the compact array containing only tigRNAs, along with the corresponding donor templates, was commercially synthesized (SynbioB, Tianjin, China) and assembled into the pCRCT-ParTasR plasmid by Gibson Assembly. The tRNA sequences used in this study are listed in **Supplementary Table 4**, and the DNA sequences of the repair templates are provided in **Supplementary Table 5**.

### Yeast transformation

Plasmid transformations of BY4741 or its derivative strains were performed using the LiAc/SS carrier DNA/PEG method^46^. For each transformation, 1 μg of plasmid DNA was used. After transformation, cells were recovered in 1 mL of YPAD at 30 ℃ with shaking at 250 rpm for 1 h, washed once with sterile water, and then processed according to the editing strategy as follows. For single locus editing, transformed cells were transferred to 2 mL of appropriate synthetic complete (SC) selective medium and cultivated at 30 ℃ with shaking at 250 rpm for 3 days, before being plated onto selective plates. For multiplex editing, transformed cells were first plated onto selective plates to obtain transformants. Colonies were then scraped from the plates, resuspended, and further cultivated in 2 mL of appropriate SC selective medium at 30 °C with shaking at 250 rpm for 3 days, before being re-plated onto selective plates for final colony examination.

### Calculation of single-gene editing efficiencies

For the SSA-based cleavage assay using the *ADE2* reporter strain, equal OD of transformed cell cultures were plated onto SC-U plates and SC-U-Adenine plates. The editing efficiency was calculated as the ratio of colony counts on SC-U-Adenine plates to those on SC-U plates. For *ADE2* editing, editing efficiency was determined by calculating the percentage of pink colonies among total colonies on SC-U plates. For *CAN1* and *LYP1* editing, 20 colonies were randomly selected from SC-U plates and replica-plated onto SC-arginine plates containing 60 μg/mL L-canavanine (for *CAN1* disruption) or SC-lysine plates containing 250 μg/mL thialysine (for *LYP1* disruption). Plates were incubated at 30 °C for 2–3 days, and editing efficiency was calculated as the percentage of colonies that grew on selective plates.

### Calculation of multiplex gene editing efficiencies

For dual-locus editing at *ADE2*, twenty pink transformants were randomly selected. Genomic DNA extracts of individual transformants were prepared using the MightyPrep reagent for DNA (Cat. # 9182, Takara Biotechnology, Hangzhou, China). The target region was PCR amplified using appropriate primers (**Supplementary Table 1**) and Sanger sequenced (Shangya Biotechnology, Hangzhou, China). The editing efficiency was calculated as the percentage of mutated sequences at both target sites. For triple-gene editing, the proportion of pink colonies was first measured to determine *ADE2* editing efficiency. Twenty pink colonies were then randomly selected and replica-plated onto SC-arginine plates containing 60 μg/mL L-canavanine and SC-lysine plates containing 250 μg/mL thialysine to assess *CAN1* and *LYP1* disruption, respectively. Colonies that remained pink on SC-U plates and grew on both canavanine and thialysine plates were regarded as triple-gene edited colonies. Selected colonies were further confirmed by Sanger sequencing of the three editing loci.

### Integration of EGFP and the lycopene pathway

For EGFP integration, four highly active *ADE2*-targeting tigRNAs were used to direct ParTasR cleavage, and a linear donor fragment containing the TDH3p-*EGFP*-ADH1t expression cassette flanked by 50 bp homology arms was co-delivered. For each transformation, 1 μg plasmid (expressing ParTasR and the tigRNA) was mixed with 1 μg of the linear donor fragment. After transformation, the proportion of EGFP-positive cells was quantified by flow cytometry. For lycopene pathway integration, a highly active *CAN1*-targeting tigRNA was used, and three linear donor fragments containing ENO2p-*crtE*-ENO2t, TEF2p-*crtYB*-TEF2t, and CDC19p-*crtI*-CDC19t, respectively, were co-delivered. The three fragments were designed with 150 bp overlapping regions between adjacent fragments to enable seamless assembly via homologous recombination in yeast. The entire pathway was flanked by 50 bp homology arms targeting the *CAN1* locus. For each transformation, 1 μg of the plasmid was mixed with 1 μg of each linear donor fragment (3 μg in total). Successful integration was confirmed by red colony phenotype and colony PCR of the junction regions.

### Mismatch tolerance assay

All tigRNA mismatches were introduced via inverse PCR using the two perfect-match plasmids as templates. The original repair templates introducing the TAA stop codon were retained in all constructs. Yeast transformation and selection were performed as described above. Briefly, for each transformation, 1 μg of plasmid DNA was used. After transformation, cells were transferred to 2 mL of SC-U selective medium and cultivated at 30 ℃ with shaking at 250 rpm for 3–4 days, then plated onto SC-U plates and incubated at 30 ℃ for an additional 2–4 days. Editing efficiency was calculated as the percentage of pink colonies among total colonies.

## SUPPLEMENTARY DATA

Supplementary data is available: Supplementary Figures 1–6 and Supplementary Tables 1–5.

## ACKNOWLEDGEMENTS

We thank iBioFoundry and Core Facility at ZJU-Hangzhou Global Scientific and Technological Innovation Center for their technical support.

## AUTHOR CONTRIBUTIONS

Z.B., Z.C. and J.Z. conceived of the study. Z.C. performed all the experiments and analyzed the data with assistance from Y.S., L.X., Y.C., and N.M.W.. Z.C. wrote the manuscript and made the figures. Z.B. and C.C. reviewed and edited the manuscript. All authors read and approved the final manuscript.

## DATA AVAILABILITY

All data supporting this study are available in the article and its supplementary information.

## AI USAGE

During the preparation of this work, the authors used GPT-5 for improving the language and grammar of the paper. The authors reviewed and edited the output as needed and take full responsibility for the content of the paper.

