## Supplementary Information for "Multiplex genome engineering in yeast using the TIGR-Tas system"

### SUPPLEMENTARY FIGURES

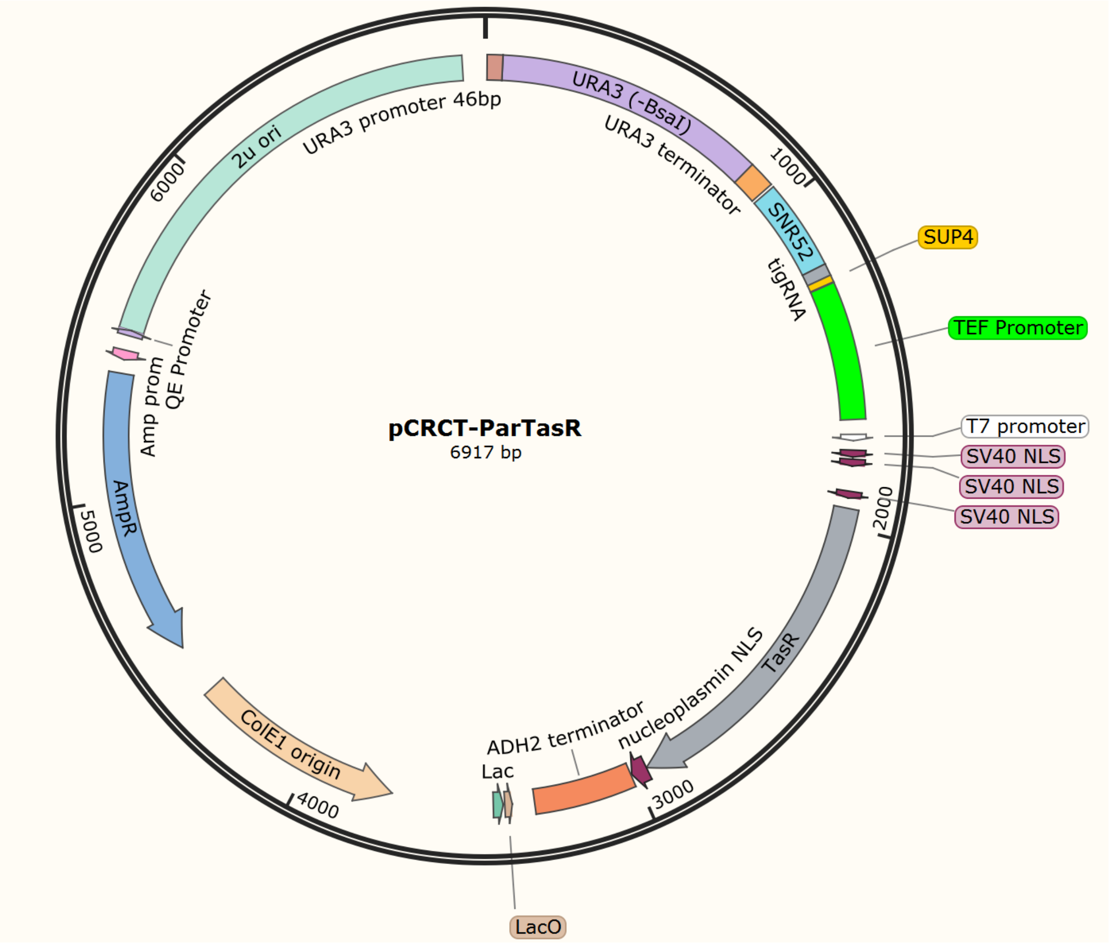


**Supplementary Figure 1. Plasmid map of pCRCT-ParTasR used in this study.** Note that this representative map does not contain donors.

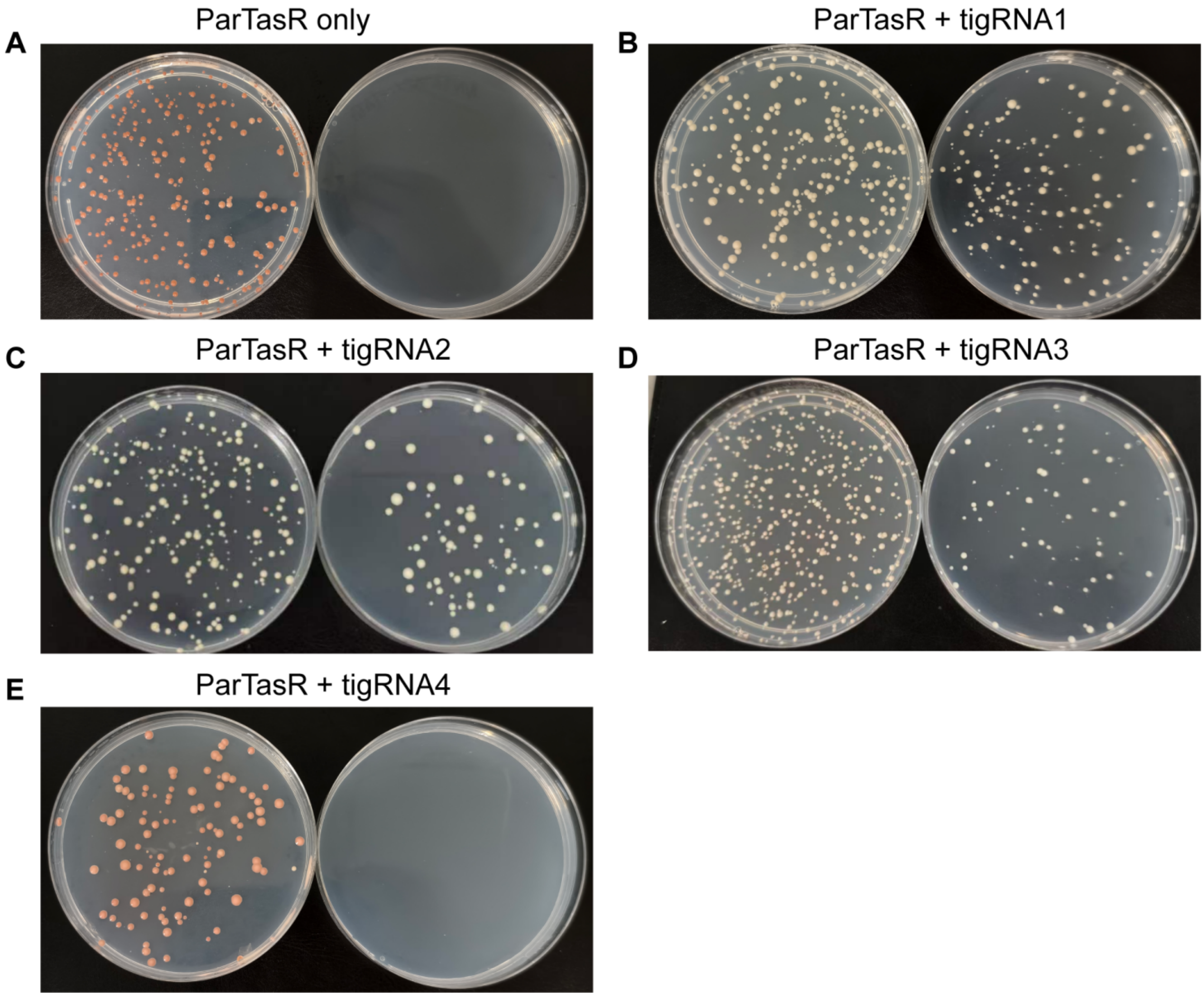


**Supplementary Figure 2. Validation of TasR DNA cleavage activity using the *ADE2* SSA reporter system. (A**) Colony phenotype of yeast cells transformed with ParTasR alone. **B–E**, Colony phenotypes of yeast cells transformed with ParTasR together with tigRNA1 (**B**), tigRNA2 (**C**), tigRNA3 (**D**), and tigRNA4 (**E**). In each panel, the left plate was supplemented with low adenine, the right plate lacked adenine. Results from one of three independent experiments were shown.

A

B

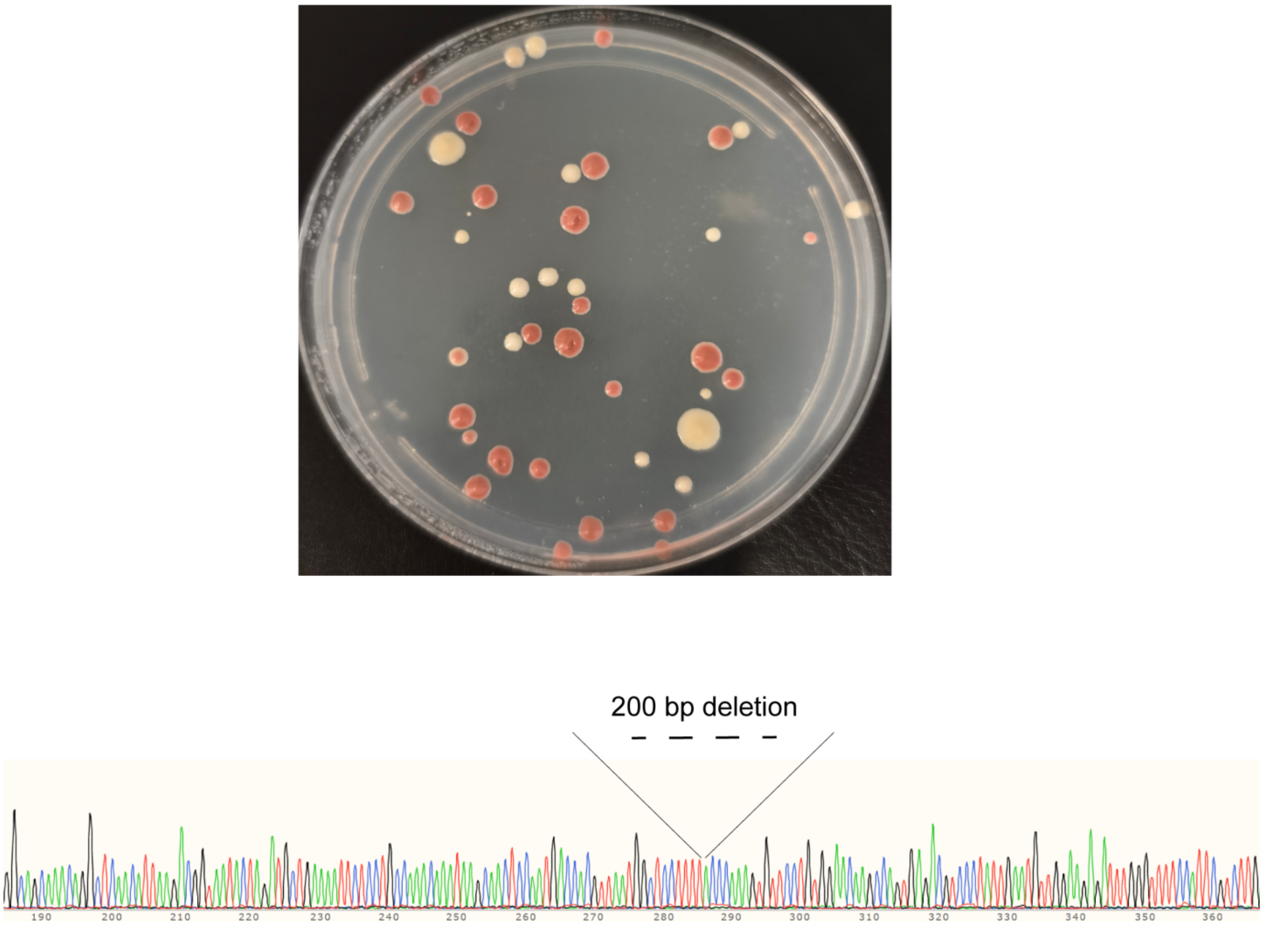


**Supplementary Figure 3. 200 bp deletion at the *ADE2* locus. (A)** Colony phenotype of yeast cells transformed with tigRNA targeting ADE2.c. **(B)** Sequencing result confirming the precise 200-bp deletion at the *ADE2* locus.

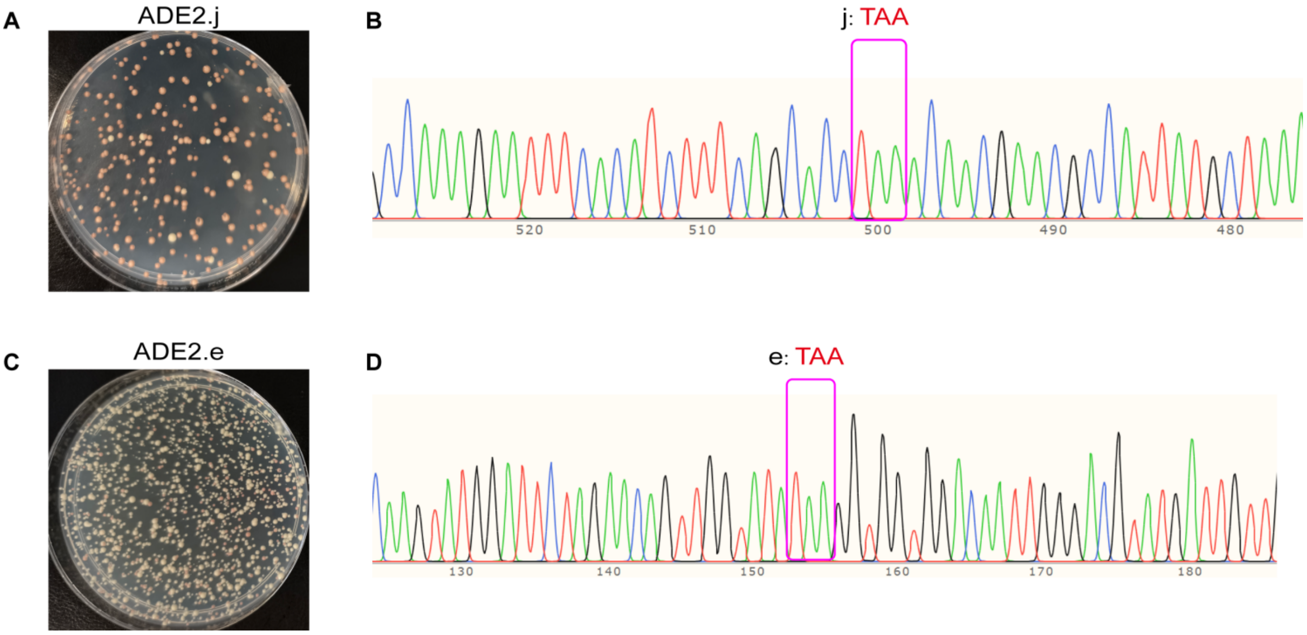


**Supplementary Figure 4. TAA stop codon introduction at *ADE2* loci.** **(A)** Colony phenotype of yeast cells transformed with tigRNA targeting ADE2.j. **(B)** Sequencing result confirming TAA stop codon introduction at the ADE2.j locus. **(C)** Colony phenotype of yeast cells transformed with tigRNA targeting ADE2.e. **(D)** Sequencing result confirming TAA stop codon introduction at the ADE2.e locus.

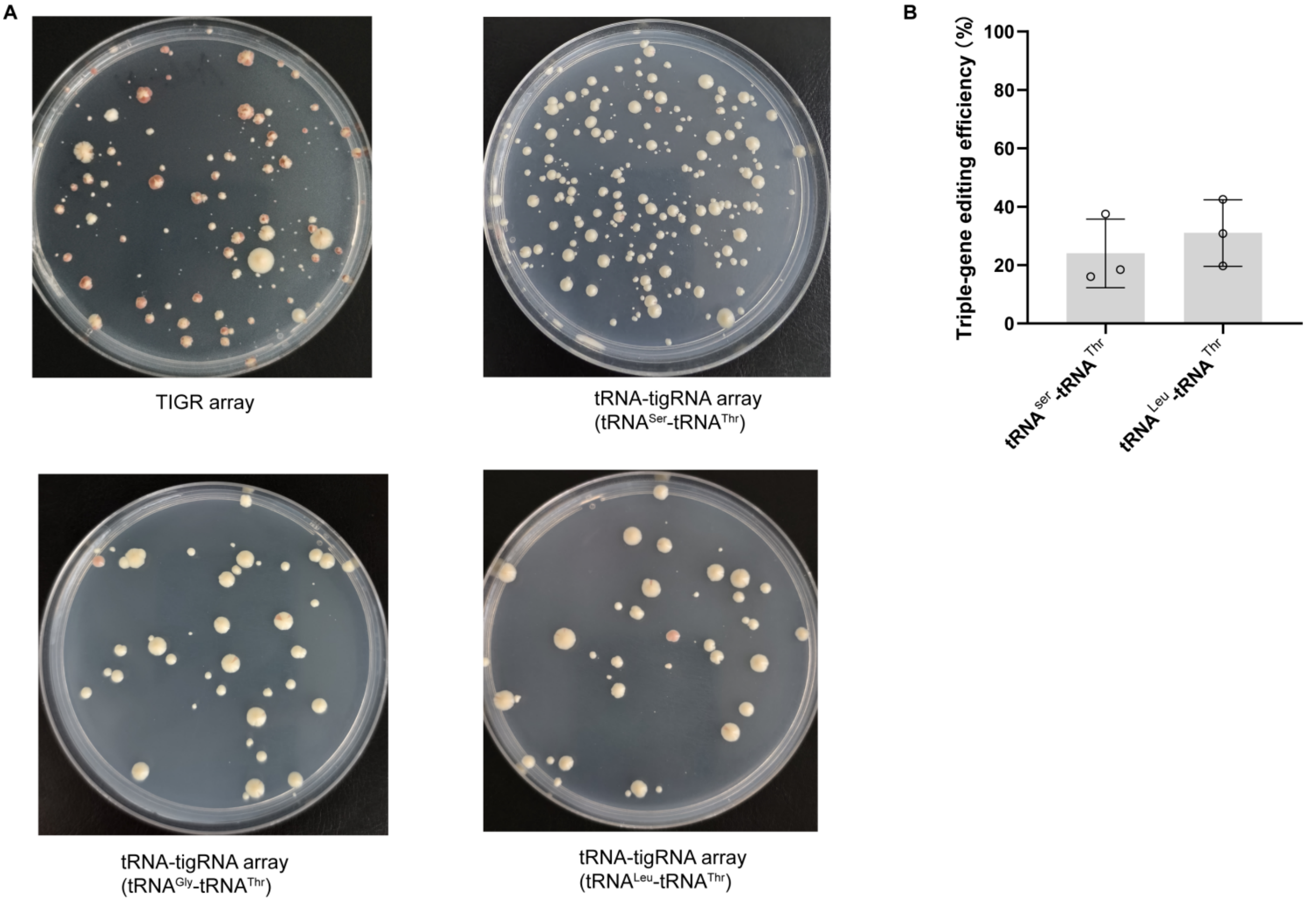


**Supplementary Figure 5. Triple-gene editing using alternative tRNA multiplexing strategies.** **(A)** Representative plates showing colony phenotypes of the TIGR array strategy and the tRNA-tigRNA array strategy. Plates were plated after transformation without further liquid culture. **(B)** Triple-gene editing efficiencies using $tRNA^{Ser}$-$tRNA^{Thr}$ and $tRNA^{Leu}$-$tRNA^{Thr}$. Data represent mean ± SD from three independent experiments.

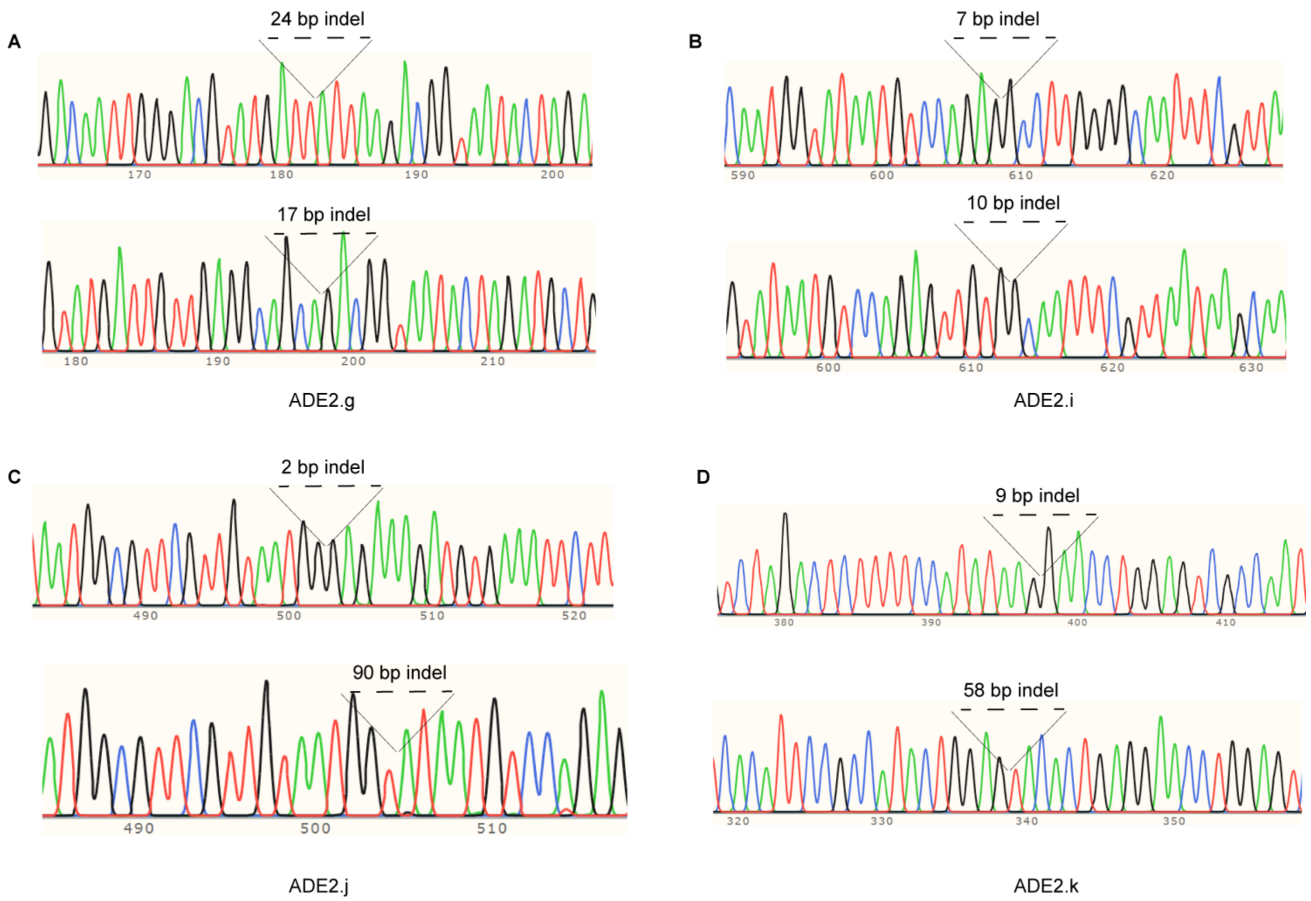


**Supplementary Figure 6. Indels generated by ParTasR cleavage in the absence of a donor template. (A)** Two representative Sanger sequencing chromatograms of the ADE2.g target site showing indels. **(B)** Two representative Sanger sequencing chromatograms of the ADE2.i target site showing indels. **(C)** Two representative Sanger sequencing chromatograms of the ADE2.j target site showing indels. **(D)** Two representative Sanger sequencing chromatograms of the ADE2.k target site showing indels.

### SUPPLEMENTARY TABLES

Supplementary Tables are supplied as separate excel sheets:

**Supplementary Table 1.** Amino acid sequence of ParTasR.

**Supplementary Table 2.** Primers used in this study.

**Supplementary Table 3.** Type III promoter sequences used to express tigRNAs.

**Supplementary Table 4.** Sequences of tRNAs used in this study.

**Supplementary Table 5.** DNA sequences of the repair templates used in this study.
